## Supplementary Text for "ADP is the dominant controller of AMP-activated protein kinase activity dynamics in skeletal muscle during exercise"

**Contents:**

1. Supplementary Methods
    a. Model construction: System definition
    b. Model construction: Topology
    c. Model calibration
    d. MPSA: Criteria for acceptable models
    e. MPSA: Framework for analyzing AMPK activation
2. Supplementary Tables S1-S7
    a. Table S1. Model species and initial concentrations
    b. Table S2. Rate equations
    c. Table S3. System of ordinary differential equations
    d. Table S4. Model parameter values
    e. Table S5. Allosteric activation potencies for the various simulations
    f. Table S6. Simulation-specific parameter values and initial conditions
    g. Table S7. Activator properties
3. Supplementary Figures
    a. Figure S1. Boxplots for the Unconstrained MPSA.
    b. Figure S2. Boxplots for the α1β2γ1 $K_D$ MPSA.

c. Figure S3. Boxplots for the α2β2γ3 $K_D$ MPSA.
d. Figure S4. Model-predicted time courses of AXP-p-AMPK in the remaining 48 models.
**4.** Supplementary References

---

### Supplementary Methods

*Model construction: System definition*

We defined our system as a single muscle fiber contracting within a primary agonist muscle during cycling exercise. Human muscle is composed of three main fiber types that differ according to the predominant myosin heavy-chain isoforms they express (type I, IIa, and IIx). The fiber types differ with respect to their speeds of contraction (slow- or fast-twitch), their metabolism (oxidative or glycolytic), and their fatiguability (fatigue-resistant, fatigable) [1]. The recruitment of the fibers depends on the task, such that we had to consider the fiber type in our study. We assumed that our muscle fiber features average contractile, metabolic, and fatigability properties in the spectrum from slow to fast-twitch fibers, and thus would most closely resemble a type-IIa fiber [2]. This assumption had two benefits. First, it allowed us to specify relatively simple recruitment profiles in response to distinct exercise modalities. In our study, we simulated two distinct types of exercise, namely continuous submaximal-intensity exercise and sprint-interval exercise (SIE), the latter of which involves repeated maximum-intensity 30-s sprints interspersed with 4-min-long passive recovery bouts. Type IIa fibers are recruited in both types of exercise [3–5], and we thus assumed that the fiber was recruited throughout the duration of exercise in a manner corresponding to the measured power profiles. Second, the assumption also enabled the valid comparison of model outputs to data from homogenates of human muscle biopsies. Biopsies contain a mixture of muscle fiber types and therefore the data represent the average response of the fibers within the biopsy. Additional simplifying assumptions included that the muscle fiber behaved as a well-stirred tank reactor and the modeled molecules were assumed to be present in sufficient concentrations so as to behave deterministically. These assumptions enabled the use of ordinary differential equations (ODEs) as the model's mathematical framework.

*Model construction: Topology*

Our model features three modules, two of which represent biochemical reactions intrinsic to the cell (*the bioenergetic module* and the *AMPK regulatory module*), as well as a *pharmacological-activator (PA)* *module*, which represents reactions involving exogenously added small-molecule activators of AMPK (Fig. 1 in the main text). The bioenergetic, AMPK regulatory, and PA modules respectively feature five, eight, and four molecular species (Table S1). These species are interconverted through intermolecular association and enzyme-catalyzed biochemical reactions, the rate equations for which are listed in Table S2. These rate equations were used within the 17 coupled nonlinear ODEs that describe the rates of change of the concentrations of the molecular species (Table S3). The species concentrations are given by sum of the reactions that generate and consume each species, expressed mathematically as follows:

$$\frac{d}{dt}[species] = \sum rates_{generation} - \sum rates_{consumption} \quad (1)$$

The rate equations associated with the bioenergetic and AMPK-regulatory modules feature 47 kinetic parameters while the PA module features ten additional parameters (Table S4). The initial concentrations for each species in the model are listed in Table S1. We converted concentrations reported in units of mmol/kg dry weight of muscle to mmol/L by multiplying by 3.13 L/kg [6].

*The bioenergetic module*. The bioenergetic module combines the previously validated model of Vicini et al. [7] with elements of the model of Lambeth et al. [8], both of which simulate AXP kinetics in contracting skeletal muscle. The module includes ATP hydrolysis (Fig. 1 and Table S2, reaction r1) and ATP supply processes including oxidative phosphorylation (Fig. 1 and Table S2, reaction r2), the creatine kinase reaction (CK; Fig. 1 and Table S2, reactions r3 and r4), and the adenylate kinase reaction (AK; Fig. 1 and Table S2, reaction r5). The rate equations and pre-calibration initial conditions for these reactions were obtained from the source models [7,8]. To foster model parsimony, we assumed a constant total adenine nucleotide pool.

Exercise was simulated by increasing the rate of ATP hydrolysis. Specifically, we changed the value of the rate constant in reaction r1 (Table S2) from a value reflecting the rate at rest ($k_{rest}$, parameter 12 in Table S4) to one commensurate with exercise ("stimulated", $k_{stim}$, parameter 13 in Table S4). Simulated exercise was ceased by setting the rate constant of the ATP hydrolysis reaction to $k_{post}$ (parameter 14 in Table S4) [7]. We simulated exercise protocols of different intensities by adjusting $k_{stim}$ until the simulated PCr concentrations matched the values measured in muscle biopsies, as PCr concentration is a well-established marker of muscle fiber recruitment and exercise intensity [9–11].

*The AMPK regulatory module*. The AMPK regulatory module featured the putative mechanisms of AXP control of AMPK activity (Fig. 1). The AMPK regulatory module consisted of eight species: unmodified and phosphorylated AMPK that were either unbound or bound to one of AMP, ADP, or ATP (Fig. 1). Our naming convention is as follows: complexes are written as AXP-(p)-AMPK, wherein the AXP is one of AMP, ADP, or ATP, the "p" indicates phosphorylation, and the suffix denotes the AMPK protein. For example, *AMP-bound phospho-AMPK* is written as *AMP-p-AMPK*, whereas *ADP bound to unphosphorylated AMPK* is written as *ADP-AMPK*.

We made the following simplifying assumptions to foster model parsimony: 1) cells contain only a single isoform of AMPK, 2) AMPK reversibly binds to the AXP at a single binding site (Table S2, reactions r6 to r11), 3) AMPK features a single activating phosphorylation site (corresponding to the Thr-172 residue), and 4) the kinetics of AMPK phosphorylation and dephosphorylation could be adequately described using Michaelis-Menten kinetics (Table S2, reactions r12 to r19).

These assumptions were justified as follows. First, while isoform-specific responses to exercise exist [12], each isoform is activated during exercise [13], such that we considered aggregate AMPK activity as the key factor driving training adaptations. Second, AXP-(p)-AMPK binding kinetics are well represented by a first-order process indicating a single binding site [14,15], which is thought to be a high-affinity AXP-binding site at the CBS3 (cystathionine-β-synthase) motif on AMPK [16]. Third, the primary mechanism of AMPK activation in mammalian cells is its phosphorylation at the Thr-172 residue [17,18]. The assumption of Michaelis-Menten kinetics was justified as follows. AMPK phosphorylation and dephosphorylation reactions each involve two reactants and an enzyme: AMPK phosphorylation involves the kinase (LKB1 or CaMKKβ), [(AXP)-AMPK], and ATP while phospho-AMPK dephosphorylation involves a phosphatase (PP1, PP2A, or PP2C [13]), (AXP)-p-AMPK, and $H_2O$. We assume that ATP and $H_2O$ concentrations are relatively constant and well in excess of the other reactants. We assumed that the kinase and phosphatase concentrations remain constant due to the transition complexes existing only for very short times, and that they are constitutively active, as has been demonstrated for LKB1 [19]. The effects of AXP binding on the AMPK phosphorylation and dephosphorylation reaction rates were modeled by using different $V_{max}$ values for each of the AXP-(p)-AMPK species (Table S4, parameters 40-47).

We quantified AMPK complex activities as $V_{max}$ values, which were calculated as the product of complex-specific $k_{cat}$ and concentration values, the latter of which varied dynamically over time. We assumed that only phosphorylated AMPK complexes were active [17,18]. In addition, AMPK is allosterically activated by AMP, which we modeled by increasing the $k_{cat}$ for AMP-p-AMPK complex relative to the $k_{cat}$ values of the other complexes (Table S5).

With respect to initial conditions, we set the total AMPK concentration equal to 0.6 mM. This choice was justified by the observations that the AMPK subunit β1 concentration is 0.06 mM in rat extensor digitorum longus (EDL) muscle, and is present in ~10% of the AMPK molecules, with the balance featuring the β2 isoform [20,21]. This total AMPK concentration was then assumed to be composed of equal concentrations of the six AXP-(p)-AMPK complexes, which resulted in their pre-calibration initial conditions (Table S1).

*The pharmacological-activator module*. The PA module adds the influence of pharmacological agents that directly bind to AMPK. Here, we modeled the activation of AMPK by the small-molecule activators 5-aminoimidazole-4-carboxamide ribonucleoside (AICAR) and Compound 991 (C991). We set the topology of the module to parsimoniously represent the experiments from published studies of AMPK activation in response to AICAR and C991 treatment. In the case of AICAR, the experiment involved perfusion of rat gastrocnemius muscle with AICAR, which is taken up by the cells and phosphorylated into the AMP analogue AICAR 5'-monophosphate (ZMP) by the enzyme adenosine kinase [22]. We introduced reactions representing the interconversion of AICAR to ZMP and the degradation of ZMP (Fig. 1 and Table S2, reactions r24 and r25). We then added the reversible binding reaction of ZMP with AMPK and p-AMPK (Fig. 1 and Table S2, r22 and r23).

The experiment with C991 involved adding bolus doses of C991 ($10^{-2}$ mM, $10^{-3}$ mM, or $10^{-4}$ mM) to the culture media overlying C2C12 murine myotube cells [23]. The action of C991 on AMPK was modeled as a binding reaction (Fig. 1 and Table S2, reactions r22 and r23). We made the simplifying assumption that both ZMP and C991 bind competitively to the AXP-binding site on AMPK, which is only true for ZMP because C991 actually binds to the allosteric drug and metabolite-binding (ADaM) site located at the interface of the kinase- and glycogen-binding domains of the α- and β-subunits, respectively [24]. We considered only β2-containing AMPK isoforms as the active complexes.

*Model calibration*

The model parameter values were calibrated using the procedure outlined by Kim et al. [25]. We first located published parameter estimates, with those for the bioenergetics module obtained from the source models [7,8] (Table S4, parameters 1-19). For the AMPK regulatory module (Table S4, parameters 20-31), the kinetic parameter values for AMP and ADP binding to AMPK were set by assuming that the forward binding constants ($k_f$) equaled 1 $mM^{-1}s^{-1}$ and then calculating the reverse binding constants ($k_r$) from the product of this $k_f$ value and the $K_D$ values reported by Xiao et al. [15] (Table S4). Those for ATP were set to reflect the $K_D$ for Mg-$ATP^{2-}$ binding to AMPK [26], given that most ATP is coordinated with magnesium *in vivo*. This $K_D$ is substantially less than that of unconjugated ATP [15]. The same values were used for AXP binding to phospho-AMPK (Table S4) [15,27]. In some cases the published values could be straightforwardly used, in other cases the units required conversion using biochemical equations (Table S4) [28]. These calculations are provided in the Supplementary Spreadsheet file. These initial estimates were collectively called the "pre-calibration" parameter values.

Most of the pre-calibration parameter values were derived from *in vitro* experiments, such that the *in vivo* values might be different. We therefore sought to adjust the parameter values so that the model reproduced measurements made from exercising humans. We used the data of Stephens et al., which features time course measurements of PCr and AXP concentrations and AMPK activity in muscle biopsies of human volunteers who cycled for 30 min at moderate intensity (~63% of $\dot{V}O_{2peak}$) [29]. We also mandated that the model replicate the qualitative patterns of the phospho-AMPK time course observed during submaximal aerobic exercise. These features included the following: 1) at least 25% of total AMPK was phosphorylated at rest [30], 2) in response to the onset of moderate-intensity exercise, phospho-AMPK levels increased gradually and substantially, and 3) upon cessation of exercise, phospho-AMPK levels returned to resting levels [31,32]. To determine the parameters that most affect the phospho-AMPK kinetics, we performed a one-factor-at-a-time local sensitivity analysis. The analysis involved adjusting each parameter value and initial condition by 10% from the pre-calibration value, simulating the model, and documenting the observed change on the features of the phospho-AMPK time course. The parameters that had the highest effects on the features were then adjusted until the three criteria were satisfied.

The model was then manually adjusted to fit the activity data [29]. Kinase activities are typically expressed in absolute units specific to a peptide substrate (i.e., mol of substrate phosphorylated per unit volume per unit time), or as fold changes relative to another condition. The absolute units are specific to the *in vitro* situation and therefore do not necessarily translate directly to the *in vivo* situation. Therefore, we set the first data point equal to the $V_{max}$ computed from the model initial conditions (in units of mmol $L^{-1}$ $s^{-1}$), followed by converting the subsequent data points into absolute activities by computing the products of the initial $V_{max}$ and the reported fold-changes in activity. In this way, the numbers predicted for the AMPK activities are not necessarily "real" in an absolute sense, but they do faithfully reflect the experimentally observed changes. The resulting parameter set is referred to as the "calibrated parameter set" (Table S4).

The kinetic parameter values for reactions involving AMPK and ZMP or C991 are listed in Tables S4, S5, and S6, and were obtained from the literature or expressed as a value relative to the corresponding parameters for AMP. Simulating the effects of ZMP introduced the complication of its precursor AICAR being the experimental treatment, such that we needed to include the conversion reaction in the model. Furthermore, the dataset used for comparison was from experiments involving AICAR perfusion of rat skeletal muscle, which necessitated an uptake reaction. We modeled these reactions using unidirectional first-order rate equations and manually tuned the parameter values so that the predicted ZMP concentration kinetics matched available ZMP time-course concentration data [33]. No calibration was performed for the parameters affecting AMPK activity when simulating the activator time courses. The pharmacological module was also used for simulating the dose-responses for each of the activators (AMP, ADP, ZMP, C991), and the corresponding activator $K_D$s, potencies, and concentration ranges employed in this analysis are listed in Table S7 [27].

*Multi-parametric sensitivity analysis: Criteria for acceptable models*

Randomly generated parameter sets will lead to numerical pathologies or biologically implausible model outputs, so we specified the following qualitative criteria to classify models as acceptable:

- All concentrations were positive values throughout the duration of the simulation.
- All concentrations remained less than 500 mM, which represents a value larger than the sum of all constituent concentrations in the model.
- All model outputs were non-complex numbers.
- Input parameters did not result in matrices that were singular or badly scaled.
- The time course ran to completion (t > 2,780 s out of a 2,800 s simulation).
- The initial amount of phospho-AMPK was less than 40% of the total AMPK.
- The initial amount of phospho-AMPK was greater than 1% of the total AMPK.
- The maximum amount of phospho-AMPK increased more than 20% higher than the initial amount of phosphorylated AMPK.
- The maximum amount of phospho-AMPK during stimulation was greater than $1/6^{th}$ of the total amount of AMPK.
- The final amount of phospho-AMPK was greater than the resting amount of AMPK.
- Phospho-AMPK levels were at steady state prior to the start of exercise [changed by less than 0.001 mM from 30 s before stimulation].

*Multi-parametric sensitivity analysis: Framework for analyzing AMPK activation*

We expect that interactions between parameters determine the dominance of a given AXP in controlling AMPK activity, but the number of parameters precludes straightforward analysis. We therefore proposed a framework for the sensitivity analysis based on simplified theoretical considerations. We first considered a reaction system containing phospho-AMPK and an allosteric activator of AMPK. At steady state, the concentration of activator-bound phospho-AMPK can be expressed as follows:

$$[act \cdot p\text{-}AMPK] = \frac{[act][p\text{-}AMPK]}{K_{D,actpAMPK}} \quad \text{(S1)}$$

where $[act \cdot p\text{-}AMPK]$ is the concentration of activator-bound phospho-AMPK, $K_{D,actpAMPK}$ is the dissociation constant for activator-bound phospho-AMPK, $[act]$ is the concentration of activator, and $[p\text{-}AMPK]$ is the concentration of phospho-AMPK. Note that the concentrations are those observed after steady state has established, not the concentrations that were added to the system at the start of the reaction.

We next defined $\alpha_{\Sigma}$, the total AMPK activity of the system if a substrate were present in saturating amounts, as the following sum:

$$\alpha_{\Sigma} = V_{actpAMPK}[act \cdot p\text{-}AMPK] + V_{pAMPK}[p\text{-}AMPK] \quad \text{(S2)}$$

in which $V_{actpAMPK}$ and $V_{pAMPK}$ are the $V_{max}$ of $act \cdot p\text{-}AMPK$ and $p\text{-}AMPK$ respectively. Substituting Equation S1 into Equation S2 gives

$$\alpha_{\Sigma} = V_{actpAMPK}\frac{[act][p\text{-}AMPK]}{K_{D,actpAMPK}} + V_{pAMPK}[p\text{-}AMPK] \quad \text{(S3)}$$

which upon simplification gives

$$\alpha_{\Sigma} = [p\text{-}AMPK]\left(V_{actpAMPK}\frac{[act]}{K_{D,actpAMPK}} + V_{pAMPK}\right) \quad \text{(S4)}$$

Phosphorylation and allostery activate AMPK in a multiplicative manner [34], such that $V_{actpAMPK}$ can be expressed as a product of allosteric activation, $\alpha_{allo}$, and $V_{pAMPK}$. Substituting this product into Equation S4 gives

$$\alpha_{\Sigma} = [p\text{-}AMPK]\left(V_{pAMPK}\alpha_{allo}\frac{[act]}{K_{D,actpAMPK}} + V_{pAMPK}\right) \quad \text{(S5)}$$

such that

$$\alpha_{\Sigma} = V_{pAMPK}[p\text{-}AMPK]\left(\alpha_{allo}\frac{[act]}{K_{D,actpAMPK}} + 1\right) \tag{S6}$$

Equation S5 expresses the five factors that determine an activator's potency for activating AMPK, i.e., the concentration of activator, the binding affinity of the activator for AMPK (expressed as $K_D$), the concentration of phospho-AMPK and its activity, and the allosteric activation of AMPK provided by the activator.

One way to compare the relative potencies of two activators (denoted by the subscripts 1 and 2) is to express as a ratio the equations for the two activators:

$$\frac{\alpha_{\Sigma,1}}{\alpha_{\Sigma,2}} = \frac{V_{pAMPK}[p\text{-}AMPK]_1\left(\alpha_{allo,1}\frac{[act_1]}{K_{D1}} + 1\right)}{V_{pAMPK}[p\text{-}AMPK]_2\left(\alpha_{allo,2}\frac{[act_2]}{K_{D2}} + 1\right)} \tag{S7}$$

Since $V_{pAMPK}$ is the same regardless of the activator, the final form is

$$\frac{\alpha_{\Sigma,1}}{\alpha_{\Sigma,2}} = \frac{[p\text{-}AMPK]_1\left(\alpha_{allo,1}\frac{[act_1]}{K_{D1}} + 1\right)}{[p\text{-}AMPK]_2\left(\alpha_{allo,2}\frac{[act_2]}{K_{D2}} + 1\right)} \tag{S8}$$

The benefits of Equation S7 is that it compactly expresses four factors that are each determined by individual parameters or intuitive groupings from the AMPK model. Specifically, $\alpha_{allo}$ is specified as an independent parameter in the model (Table S5), the $k_f$ and $k_r$ forward and reverse rate constants for AXP binding to AMPK and phospho-AMPK determine the $K_D$, while $[act]$ is determined primarily by the bioenergetic reactions. Furthermore, computing compound ratios of these parameters or groupings of parameters that pertain to different activators should provide insights into their sensitivities.

While useful for organizing our sensitivity analysis, the above equations *cannot* be directly used to validly estimate AMPK activities. In a real system containing AMPK kinases and phosphatases, the determination of $[p\text{-}AMPK]$ is complex because its levels are determined by the activators and their propensities to enhance upstream kinase activity and reduce phosphatase activity towards AMPK. Accordingly, $[p\text{-}AMPK]$ depends on the enzyme kinetic parameters of AMPK kinases and phosphatases, as well as on $[act]$ and the $K_D$. In addition, the presence of ATP in the system causes further complexities because it is both necessary for kinase activity (as a substrate) but also competes with AMP and ADP for binding. The complexities of the real system make it difficult to express AMPK activity control as a tractable analytical equation and emphasizes the need for kinetic models to study it.

### Supplementary Tables

**Table S1**. Model species and initial concentrations. Model species (state variables) and their initial concentrations before and after the model calibration.

| **Species** | **Name** | **Pre-calibration Value (mM)** | **Calibrated Value (mM)** | **Basis for values** | | | |
|---|---|---|---|---|---|---|---|
| | | | | Ref | Calc | Calib | Est |
| | | *Bioenergetic module* | | | | | |
| $x(1)$ | ATP | 8.2 [7] | 7.66 | ☒ | ☐ | ☒ | ☐ |
| $x(2)$ | ADP | $1.3\times10^{-2}$ [7] | $5.68\times10^{-2}$ | ☒ | ☐ | ☒ | ☐ |
| $x(3)$ | AMP | $2.0\times10^{-5}$ [7] | $4.44\times10^{-4}$ | ☒ | ☐ | ☒ | ☐ |
| $x(4)$ | PCr | 32.1 [7] | 22.1 | ☒ | ☐ | ☒ | ☐ |
| $x(5)$ | $P_i$ | 3.183 [7] | 3.183 | ☒ | ☐ | ☐ | ☐ |
| | | *AMPK regulatory module* | | | | | |
| $x(6)$ | ATP-AMPK* | $1\times10^{-1}$ [21,35] | 0.075 | ☒ | ☒ | ☒ | ☐ |
| $x(7)$ | ADP-AMPK* | $1\times10^{-1}$ [21,35] | 0.075 | ☒ | ☒ | ☒ | ☐ |
| $x(8)$ | AMP-AMPK* | $1\times10^{-1}$ [21,35] | 0.075 | ☒ | ☒ | ☒ | ☐ |
| $x(9)$ | ATP-p-AMPK* | $1\times10^{-1}$ [21,35] | 0.075 | ☒ | ☒ | ☒ | ☐ |
| $x(10)$ | ADP-p-AMPK* | $1\times10^{-1}$ [21,35] | 0.075 | ☒ | ☒ | ☒ | ☐ |
| $x(11)$ | AMP-p-AMPK* | $1\times10^{-1}$ [21,35] | 0.075 | ☒ | ☒ | ☒ | ☐ |
| $x(12)$ | AMPK* | 0 | 0.075 | ☒ | ☒ | ☒ | ☐ |
| $x(13)$ | p-AMPK* | 0 | 0.075 | ☒ | ☒ | ☒ | ☐ |
| | | *Pharmacological activator module*** | | | | | |
| $x(14)$ | Act-AMPK | 0 | | ☒ | ☐ | ☐ | ☐ |
| $x(15)$ | Act-p-AMPK | 0 | | ☒ | ☐ | ☐ | ☐ |
| $x(16)$ | Act | 0 | | ☒ | ☐ | ☐ | ☐ |
| $x(17)$ | AICAR | 0 | | ☒ | ☐ | ☐ | ☐ |

Ref, references; Calc, calculations; Calib, calibrations; Est, estimations.

* Total AMPK was set to 0.6 mM and divided equally between the six nucleotide-bound complexes prior to calibration. We found that the initial concentration had no effect on outcomes, and so we divided all AMPK complexes equally post-calibration.

**The initial parameter values were based on ZMP, values for Compound 991 are presented in Table S6.

**Table S2.** Rate equations.

| Description | Equation |
|---|---|
| Rate of ATP hydrolysis | $r1 = k_{stim} \times x(1)$ or $r1 = k_{rest} \times x(1)$ |
| Oxidative Phosphorylation | $r2 = \dfrac{V_{maxOxPhos} \times \left(\frac{x(2)}{K_{ADP}}\right)^{nH}}{1 + \left(\frac{x(2)}{K_{ADP}}\right)^{nH}}$ |
| Forward Creatine Kinase | $r3 = \dfrac{\frac{V_{forCK} \times x(2) \times x(4)}{K_{ia} \times K_b}}{1 + \frac{x(2)}{K_{ia}} + \frac{x(4)}{K_{ib}} + \frac{x(1)}{K_{iq}} + \frac{x(2) \times x(4)}{K_{ia} \times K_b} + \frac{\left(TCr - x(4)\right) \times x(1)}{K_{iq} \times K_p}}$ |
| Reverse Creatine Kinase | $V_{revCK} = \dfrac{V_{forCK} \times K_{iq} \times K_p}{K_{eqCK} \times K_{ia} \times K_b}$ <br> $r4 = \dfrac{\frac{V_{revCK} \times x(1) \times \left(TCr - x(4)\right)}{K_{iq} \times K_p}}{1 + r4 + \left(\frac{x(2)}{K_{ia}}\right) + \left(\frac{x(4)}{K_{ib}}\right) + \left(\frac{x(1)}{K_{iq}}\right) + \frac{x(2) \times x(4)}{K_{ia} \times K_b} + \frac{\left(\left(TCr - x(4)\right) \times x(1)\right)}{K_{iq} \times K_p}}$ |
| Forward Adenylate Kinase | $r5f = \dfrac{\frac{V_{forAK} \times x(1)}{kmt * kmm}}{1 + \frac{x(1)}{kmt} + \frac{x(3)}{kmm} + \frac{x(1) \times x(3)}{kmt \times kmm} + \frac{2 * x(2)}{kmd} + \frac{\left(x(2)\right)^2}{kmd^2}}$ |
| Reverse Adenylate Kinase | $V_{revAK} = \dfrac{V_{forAK} \times kmd^2}{kmt \times K_{eqAK} \times kmm}$ <br> $r5r = \dfrac{\frac{V_{revAK} \times \left(x(2)\right)^2}{kmd^2}}{1 + \frac{x(1)}{kmt} + \frac{x(3)}{kmm} + \frac{x(1) \times x(3)}{kmt \times kmm} + \frac{2 * x(2)}{kmd} + \frac{\left(x(2)\right)^2}{kmd^2}}$ |
| AMPK binding ATP | $r6 = k6f \times x(1) \times x(12) - k6r \times x(6)$ |
| AMPK binding ADP | $r7 = k7f \times x(2) \times x(12) - k7r \times x(7)$ |
| AMPK binding AMP | $r8 = k8f \times x(3) \times x(12) - k8r \times x(8)$ |

| | |
|---|---|
| p-AMPK binding ATP | $r9 = k9f \times x(1) \times x(13) - k9r \times x(9)$ |
| p-AMPK binding ADP | $r10 = k10f \times x(2) \times x(13) - k10r \times x(10)$ |
| p-AMPK binding AMP | $r11 = k11f \times x(3) \times x(13) - k11r \times x(11)$ |
| Phosphorylation AMPK | $r12 = \frac{V_{maxAMPKK} \times x(12)}{km12 + x(12)}$ |
| Dephosphorylation p-AMPK | $r13 = \frac{V_{maxPPaseAMPK} \times x(13)}{km13 + x(13)}$ |
| Phosphorylation ATP-AMPK | $r14 = \frac{V_{maxAMPKKT} \times x(6)}{km14 + x(6)}$ |
| Dephosphorylation ATP-p-AMPK | $r15 = \frac{V_{maxPPaseATP} \times x(9)}{km15 + x(9)}$ |
| Phosphorylation ADP-AMPK | $r16 = \frac{V_{maxAMPKKD} \times x(7)}{km16 + x(7)}$ |
| Dephosphorylation ADP-p-AMPK | $r17 = \frac{V_{maxPPaseADP} \times x(10)}{km17 + x(10)}$ |
| Phosphorylation AMP-AMPK | $r18 = \frac{V_{maxAMPKKM} \times x(8)}{km18 + x(8)}$ |
| Dephosphorylation AMP-p-AMPK | $r19 = \frac{V_{maxPPaseAMP} \times x(11)}{km19 + x(11)}$ |
| Phosphorylation Activator-AMPK | $r20 = V_{maxAMPKKZMP} \times x(14)$ |
| Dephosphorylation Activator-p-AMPK | $r21 = V_{maxPPaseZMP} \times x(15)$ |
| AMPK binding Activator | $r22 = k12f \times x(12) \times x(16) - k12r \times x(14)$ |

| | |
|---|---|
| p-AMPK binding Activator | $r23 = k13f \times x(13) \times x(16) - k13r \times x(15)$ |
| AICAR forming Activator | $r24 = k_{AICAR} \times x(17)$ |
| Activator degradation rate | $r25 = k_{Act} \times x(16)$ |

**Table S3.** System of ordinary differential equations.

| Variable | Differential equation (with respect to time) |
|---|---|
| ATP | dx(1) = - r1 + r2 - CK - AK - r6 - r9; |
| ADP | dx(2) = + r1 - r2 + CK + 2×AK - r7 - r10; |
| AMP | dx(3) = - AK - r8 - r11 - r20; |
| PCr | dx(4) = + CK; |
| Pi | dx(5) = + r1 - r2; |
| ATP-AMPK | dx(6) = + r6 - r14 + r15; |
| ADP-AMPK | dx(7) = + r7 - r16 + r17; |
| AMP-AMPK | dx(8) = + r8 - r18 + r19; |
| ATP-p-AMPK | dx(9) = + r9 + r14 - r15; |
| ADP-p-AMPK | dx(10) = + r10 + r16 - r17; |
| AMP-p-AMPK | dx(11) = + r11 + r18 - r19; |
| AMPK | dx(12) = - r6 - r7 - r8 - r12 + r13; |
| p-AMPK | dx(13) = - r9 - r10 - r11 + r12 - r13; |
| Activator-AMPK | dx(14) = - r20 + r21 + r22; |
| Activator-p-AMPK | dx(15) = + r20 - r21 + r23; |
| Activator | dx(16) = - r22 - r23 + r24 - r25;* |
| AICAR | dx(17) = 0; |

*Activator uptake and depletion rates were set to zero when simulating dose-response experiments.

**Table S4**. Model parameter values.

| Reaction | (#) Kinetic parameter | Pre-calibration value | Calibrated value | Units | Basis for values | | | |
|---|---|---|---|---|---|---|---|---|
| | | | | | Ref* | Calc | Calib | Est |
| *Bioenergetic module* | | | | | | | | |
| Creatine kinase | (1) $V_{forCK}$ | $1.00\times10^{2}$ [7] | $1.00\times10^{2}$ | mM/s | ☒ | ☐ | ☐ | ☐ |
| | (2) $K_{b}$ | 1.11 [7] | 1.11 | mM | ☒ | ☐ | ☐ | ☐ |
| | (3) $K_{ia}$ | 0.135 [7] | 0.135 | mM | ☒ | ☐ | ☐ | ☐ |
| | (4) $K_{ib}$ | 3.9 [7] | 3.9 | mM | ☒ | ☐ | ☐ | ☐ |
| | (5) $K_{iq}$ | 3.5 [7] | 3.5 | mM | ☒ | ☐ | ☐ | ☐ |
| | (6) $K_{p}$ | 3.8 [7] | 3.8 | mM | ☒ | ☐ | ☐ | ☐ |
| | (7) $K_{eqCK}$ | $1.77\times10^{(9\text{-pH})}$ [7] | $1.77\times10^{(9\text{-pH})}$ | unitless | ☒ | ☐ | ☐ | ☐ |
| | (8) TCr | 42 [7] | 39 | mM | ☒ | ☐ | ☐ | ☐ |
| Oxidative phosphorylation | (9) $K_{ADP}$ | $5.8\times10^{-2}$ [7] | $5.8\times10^{-2}$ | mM | ☒ | ☐ | ☐ | ☐ |
| | (10) $V_{maxOxPhos}$ | 0.5 [7] | 0.5 | mM/s | ☒ | ☐ | ☐ | ☐ |
| | (11) nH | 2.568 [7] | 2.568 | unitless | ☒ | ☐ | ☐ | ☐ |
| ATP hydrolysis | (12) $k_{rest}$ | $1.4\times10^{-3}$ [7] | $2.6\times10^{-2}$ | 1/s | ☒ | ☐ | ☒ | ☐ |
| | (13) $k_{stim}$ | $1.39\times10^{-2}$ [7] | $5.0\times10^{-2}$ | 1/s | ☒ | ☐ | ☒ | ☐ |
| | (14) $k_{post}$ | NA | $2.6\times10^{-2}$ | 1/s | ☐ | ☐ | ☐ | ☒ |
| Adenylate kinase | (15) $V_{forAK}$ | 14.66 [8] | 14.66 | mM/s | ☒ | ☒ | ☐ | ☐ |

| Reaction | (#) Kinetic parameter | Pre-calibration value | Calibrated value | Units | Basis for values | | | |
|---|---|---|---|---|---|---|---|---|
| | | | | | Ref* | Calc | Calib | Est |
| | (16) kmt | 0.27 [8] | 0.27 | mM | ☒ | ☐ | ☐ | ☐ |
| | (17) kmd | 0.35 [8] | 0.35 | mM | ☒ | ☐ | ☐ | ☐ |
| | (18) kmm | 0.32 [8] | 0.32 | mM | ☒ | ☐ | ☐ | ☐ |
| | (19) $K_{eqADK}$ | 2.21 [36,37] | 0.744 | unitless | ☒ | ☐ | ☒ | ☐ |
| *AMPK regulatory module* | | | | | | | | |
| ATP-AMPK | (20) k6f | 1 [15] | 1 | 1/(mM×s) | ☒ | ☒ | ☒ | ☐ |
| | (21) k6r | $1.8\times10^{-2}$ [15] | $2.2\times10^{-1}$ | 1/s | ☒ | ☒ | ☒ | ☐ |
| ADP-AMPK | (22) k7f | 1 [15] | 1.5 | 1/(mM×s) | ☒ | ☒ | ☒ | ☐ |
| | (23) k7r | $1.5\times10^{-3}$ [15] | $2.25\times10^{-3}$ | 1/s | ☒ | ☒ | ☒ | ☐ |
| AMP-AMPK | (24) k8f | 1 [15] | 1.5 | 1/(mM×s) | ☒ | ☒ | ☒ | ☐ |
| | (25) k8r | $2.5\times10^{-3}$ [15] | $3.75\times10^{-3}$ | 1/s | ☒ | ☒ | ☒ | ☐ |
| ATP-p-AMPK | (26) k9f | 1 [15] | 1 | 1/(mM×s) | ☒ | ☒ | ☒ | ☐ |
| | (27) k9r | $1.8\times10^{-2}$ [15] | $4.0\times10^{-1}$ | 1/s | ☒ | ☒ | ☒ | ☐ |
| ADP-p-AMPK | (28) k10f | 1 [15] | 1.5 | 1/(mM×s) | ☒ | ☒ | ☒ | ☐ |
| | (29) k10r | $1.5\times10^{-3}$ [15] | $2.25\times10^{-3}$ | 1/s | ☒ | ☒ | ☒ | ☐ |
| AMP-p-AMPK | (30) k11f | 1 [15] | 1 | 1/(mM×s) | ☒ | ☒ | ☒ | ☐ |
| | (31) k11r | $2.5\times10^{-3}$ [15] | $3.75\times10^{-3}$ | 1/s | ☒ | ☒ | ☒ | ☐ |

| Reaction | (#) Kinetic parameter | Pre-calibration value | Calibrated value | Units | Basis for values | | | |
|---|---|---|---|---|---|---|---|---|
| | | | | | Ref* | Calc | Calib | Est |
| ATP-(p)-AMPK kinase and phosphatase | (32) $K_M12$ | 1.4 [38] | 1.4 | mM | ☒ | ☐ | ☒ | ☐ |
| | (33) $K_M13$ | $6.7\times10^{-2}$ [39] | $6.7\times10^{-2}$ | mM | ☒ | ☐ | ☒ | ☐ |
| (p)-AMPK kinase and phosphatase | (34) $K_M14$ | 1.4 [38] | 1.4 | mM | ☒ | ☐ | ☒ | ☐ |
| | (35) $K_M15$ | $6.7\times10^{-2}$ [39] | $6.7\times10^{-2}$ | mM | ☒ | ☐ | ☒ | ☐ |
| ADP-(p)-AMPK kinase and phosphatase | (36) $K_M16$ | 1.4 [38] | 1.4 | mM | ☒ | ☐ | ☒ | ☐ |
| | (37) $K_M17$ | $6.7\times10^{-2}$ [39] | $6.7\times10^{-2}$ | mM | ☒ | ☐ | ☒ | ☐ |
| AMP-(p)-AMPK kinase and phosphatase | (38) $K_M18$ | 1.4 [38] | 1.4 | mM | ☒ | ☐ | ☒ | ☐ |
| | (39) $K_M19$ | $6.7\times10^{-2}$ [39] | $6.7\times10^{-2}$ | mM | ☒ | ☐ | ☒ | ☐ |
| AMPK kinase $V_{max}$ | (40) $V_{maxKinase}$ | $3.92\times10^{-2}$ [38] | $5\times10^{-3}$ | mM/s | ☒ | ☒ | ☒ | ☐ |
| | (41) $V_{maxKinaseATP}$ | $3.92\times10^{-2}$ [38] | $5\times10^{-3}$ | mM/s | ☒ | ☒ | ☒ | ☐ |
| | (42) $V_{maxKinaseADP}$ | $3.92\times10^{-2}$ [38] | $7.5\times10^{-3}$ | mM/s | ☒ | ☒ | ☒ | ☐ |
| | (43) $V_{maxKinaseAMP}$ | $3.92\times10^{-2}$ [38] | $2.0\times10^{-2}$ | mM/s | ☒ | ☒ | ☒ | ☐ |
| p-AMPK phosphatase $V_{max}$ | (44) $V_{maxPPase}$ | $1.1\times10^{-1}$ [39] | $1\times10^{-2}$ | mM/s | ☒ | ☒ | ☒ | ☒ |
| | (45) $V_{maxPPaseATP}$ | $1.1\times10^{-1}$ [39] | $1\times10^{-2}$ | mM/s | ☒ | ☒ | ☒ | ☒ |
| | (46) $V_{maxPPaseADP}$ | $1.1\times10^{-1}$ [39] | $1\times10^{-3}$ | mM/s | ☒ | ☒ | ☒ | ☒ |
| | (47) $V_{maxPPaseAMP}$ | $1.1\times10^{-1}$ [39] | $1\times10^{-4}$ | mM/s | ☒ | ☒ | ☒ | ☒ |
| *Pharmacological activator module*** | | | | | | | | |

| Reaction | (#) Kinetic parameter | Pre-calibration value | Calibrated value | Units | Basis for values | | | |
|---|---|---|---|---|---|---|---|---|
| | | | | | Ref* | Calc | Calib | Est |
| ZMP-(p)-AMPK kinase and phosphatase parameters | (48) $V_{maxAMPKKZMP}$ | NA | $2.0\times10^{-2}$ | mM/s | ☐ | ☐ | ☐ | ☒ |
| | (49) $V_{maxPPaseZMP}$ | NA | $1\times10^{-4}$ | mM/s | ☐ | ☐ | ☐ | ☒ |
| | (50) $K_M20$ | NA | 1.4 | mM | ☐ | ☐ | ☐ | ☒ |
| | (51) $K_M21$ | NA | $6.7\times10^{-2}$ | mM | ☐ | ☐ | ☐ | ☒ |
| ZMP-AMPK binding | (52) k12f | NA | 1 [27] | 1/(mM×s) | ☒ | ☐ | ☐ | ☐ |
| | (53) k12r | NA | $1.8\times10^{-2}$ [27] | 1/s | ☒ | ☐ | ☐ | ☐ |
| ZMP-p-AMPK binding | (54) k13f | NA | 1 [27] | 1/(mM×s) | ☒ | ☐ | ☐ | ☐ |
| | (55) k13r | NA | $1.8\times10^{-2}$ [27] | 1/s | ☒ | ☐ | ☐ | ☐ |
| Infusion rate (AICAR to ZMP) | (56) $k_{AICAR}$ | NA | $8\times10^{-4}$ | 1/s | ☐ | ☐ | ☒ | ☐ |
| Degradation rate | (57) $k_{Act}$ | NA | $3\times10^{-3}$ | 1/s | ☐ | ☐ | ☒ | ☐ |
| *Additional parameters* | | | | | | | | |
| Muscle pH | pH | 7.0 [7,36] | 7.2 | | ☒ | ☐ | ☐ | ☐ |
| AMPK turnover number | $k_{catAMPK}$ | 30.6 [40] | 34.56 | /s | ☒ | ☒ | ☒ | ☐ |

*Ref, references; Calc, calculations; Calib, calibrations; Est, estimations.

**The initial pharmacological activator parameter values were based on ZMP, the values for Compound 991 are listed in Table S6.

**Table S5.** Allosteric activation potencies for simulations.

| Simulation set | Value | Activator(s) | Fold-activation | Reference |
|---|---|---|---|---|
| Calibrated model | Base | AMP | 4 | [24,41] |
| MPSA, α1β2γ1 | Minimum | AMP | 1 | [30,42] |
| | Base | | 4 | |
| | Maximum | | 13 | |
| MPSA, α2β2γ3 | Minimum | AMP | 1 | [26] |
| | Base | | 1.54 | |
| | Maximum | | 2 | |
| Dose-response | Base | AMP | 4 | [24,41] |
| | | ZMP | 2 | [24,41] |
| | | C991 | 3.9 | [24] |

**Table S6**. Simulation-specific parameter values and initial conditions. Summary of the changes to the parameter values and initial concentrations from the original model to simulate the exercise protocols from different studies.

| **Protocol** | **Initial concentrations** | **Kinetic parameter values** | **Reference** |
|---|---|---|---|
| Stephens | See Table S1 | See Table S4 | n/a |
| Gibala | $ATP_0$ = 9.6 mM<br>$PCr_0$ = 26.2 mM<br>$TCr_0$ = 39.9 mM | $k_{rest}$ and $k_{post}$ = $1.7\times10^{-2}$ /s<br>$k_{stim}$ = $1.05\times10^{-1}$ /s | n/a |
| ZMP | | See Table S4 | n/a |
| Compound 991 | | k12f = 1 /(mM×s)<br>k12r = $5.1\times10^{-4}$ /s<br>k13f = 1 /(mM×s)<br>k13r = $5.1\times10^{-4}$ /s<br>$V_{maxPPaseZMP}$ = $2\times10^{-4}$ mM/s<br>$k_{AICAR}$ = 0;<br>$k_{Act}$ = 0; | [24] |

**Table S7.** Activator properties for the dose-response analysis. 'Baseline' refers to the calibrated value for unphosphorylated and unbound AMPK.

| Activator Name | Observed $K_D$ | Potency | | | Concentration Range |
|---|---|---|---|---|---|
| | | *Increased Thr172 Phosphorylation* | *Decreased Thr172 Dephosphorylation* | *Allosteric Activation* | |
| Compound 991 | 0.51 µM [24] | Assumed same as AMP | C991 $V_{max} = 2 \times V_{maxPPaseAMP}$ [24] | 3.9-fold* [24] | 0.1-10 µM [23] |
| ZMP | 18 µM [27] | Assumed same as AMP [41] | ZMP $V_{max} = V_{maxPPaseAMP}$ [41] | 2-fold [41] | 500-2,000 µM [33] |
| ADP | 1.5 µM [15,26] | ⇧1.5-fold vs. baseline [42] | See Table S4 | No effect | 50-200 µM [15] |
| AMP | 2.5 µM [15,26] | ⇧4-fold vs. baseline [42] | See Table S4 | 13-fold [30,42] | 0.5-5 µM [15] |

*Average value from two AMPK isoforms, α1β2γ1 and α2β2γ1, for which data were available.

### Supplementary Figures

*Interpretation of Supplementary Figures S1 to S3*

Supplementary Figures S1 to S3 present boxplots that summarize the distributions of parameter values, ratios of these values, and composite ratios for the "ADP-dominant" (green) and "AMP-dominant" (blue) models amongst the acceptable models from the MPSAs. The fraction of ADP-mediated control of AMPK activity was calculated using Equation 2 from the main text. Models for which the fraction of ADP control exceeded 0.8 were classified as "ADP-dominant" whereas those less than 0.2 were classified as "AMP-dominant". Shifts of the blue and green boxplots relative to each other for a given parameter or ratio indicate that the quantity affects AMP or ADP dominance.

***Boxplots***. All values were plotted on the base-10 logarithmic scale. The black points represent the median, the box bounds are the first and third quartile values, the whiskers extend to the farthest point within 1.5× of the interquartile range, and the points beyond the whiskers are plotted as individual circles and are considered outliers. The horizontal red lines demarcate the bounds of the sampling ranges for the parameter values used in the MPSA. The location and width of the boxplots within the bounds indicate parameter regimes conducive to generating acceptable models.

***Variable labels***. Subscripts indicate the phosphorylation state (p = phosphorylated or 0 = dephosphorylated) and species bound to an adenine nucleotide are either preceded or succeeded by the identity of the bound nucleotide.

***Sample sizes***. The sample sizes for the MPSA boxplots were uneven because they depended on the number of acceptable models within the 50,000 simulations for each MPSA and the number of ADP- and AMP-dominant cases. The number of samples comprising each boxplot in each MPSA were as follows:

- Unconstrained: ADP, 61; AMP, 127
- α1β2γ1 $K_D$: ADP, 2,289; AMP, 18
- α2β2γ3 $K_D$: ADP, 1,390; AMP, 195

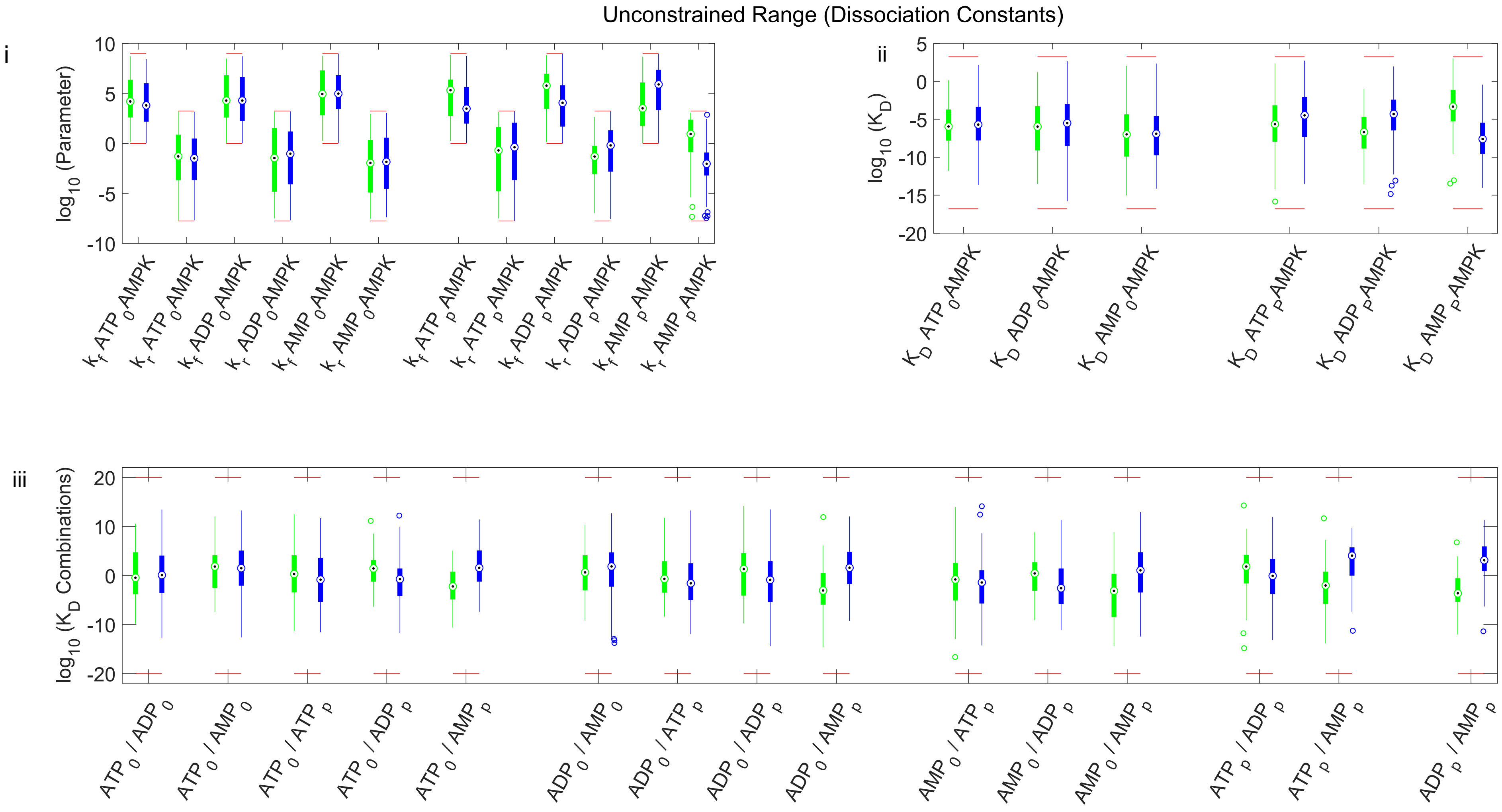


**Figure S1A**. Effects of AXP-(p)-AMPK binding constants on AMP- versus ADP-dominant control of AMPK activity in the Unconstrained MPSA. *i)* Boxplots summarize the individual parameter values representing the forward ($k_f$) and reverse ($k_r$) rate constants. *ii)* Boxplots summarize the dissociation constants ($K_D$) for each AXP-(p)-AMPK reaction, calculated as $k_r/k_f$ from each simulation. *iii)* Boxplots summarize the ratios of all possible pairs of $K_D$ values.

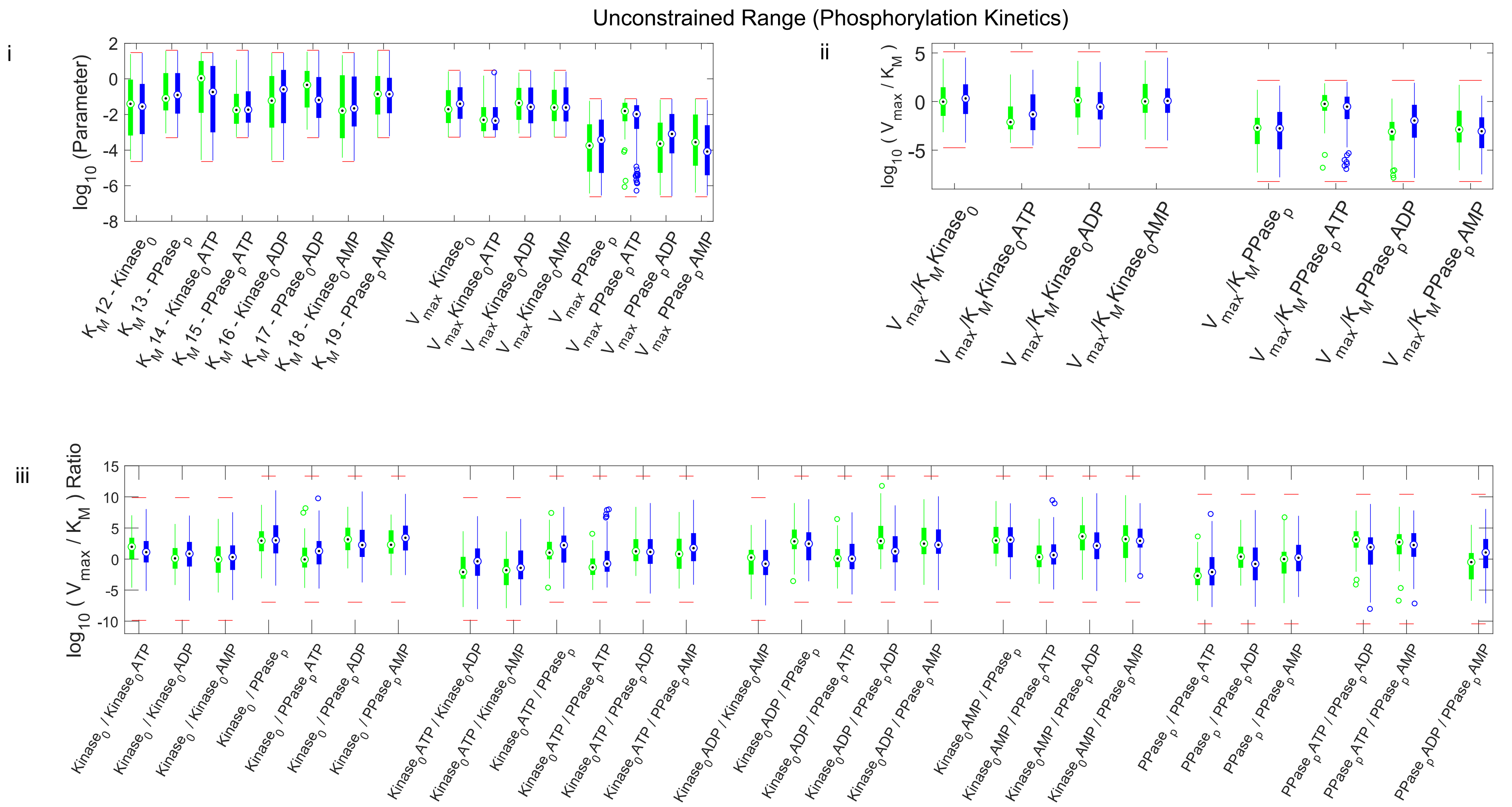


**Figure S1B.** Effects of parameters determining the enhancement of AMPK phosphorylation on AMP- versus ADP-dominant control of AMPK activity in the Unconstrained MPSA. *i)* Boxplots summarize the Michaelis constants ($K_M$) and maximum enzyme velocities ($V_{max}$) for the AMPK kinase and phosphatase. *ii)* Boxplots summarize the catalytic efficiencies of the AMPK kinase and phosphatase, which was calculated as $V_{max}/K_M$ for each simulation. *iii)* Boxplots summarize the compound ratios of kinase and phosphatase catalytic efficiencies.

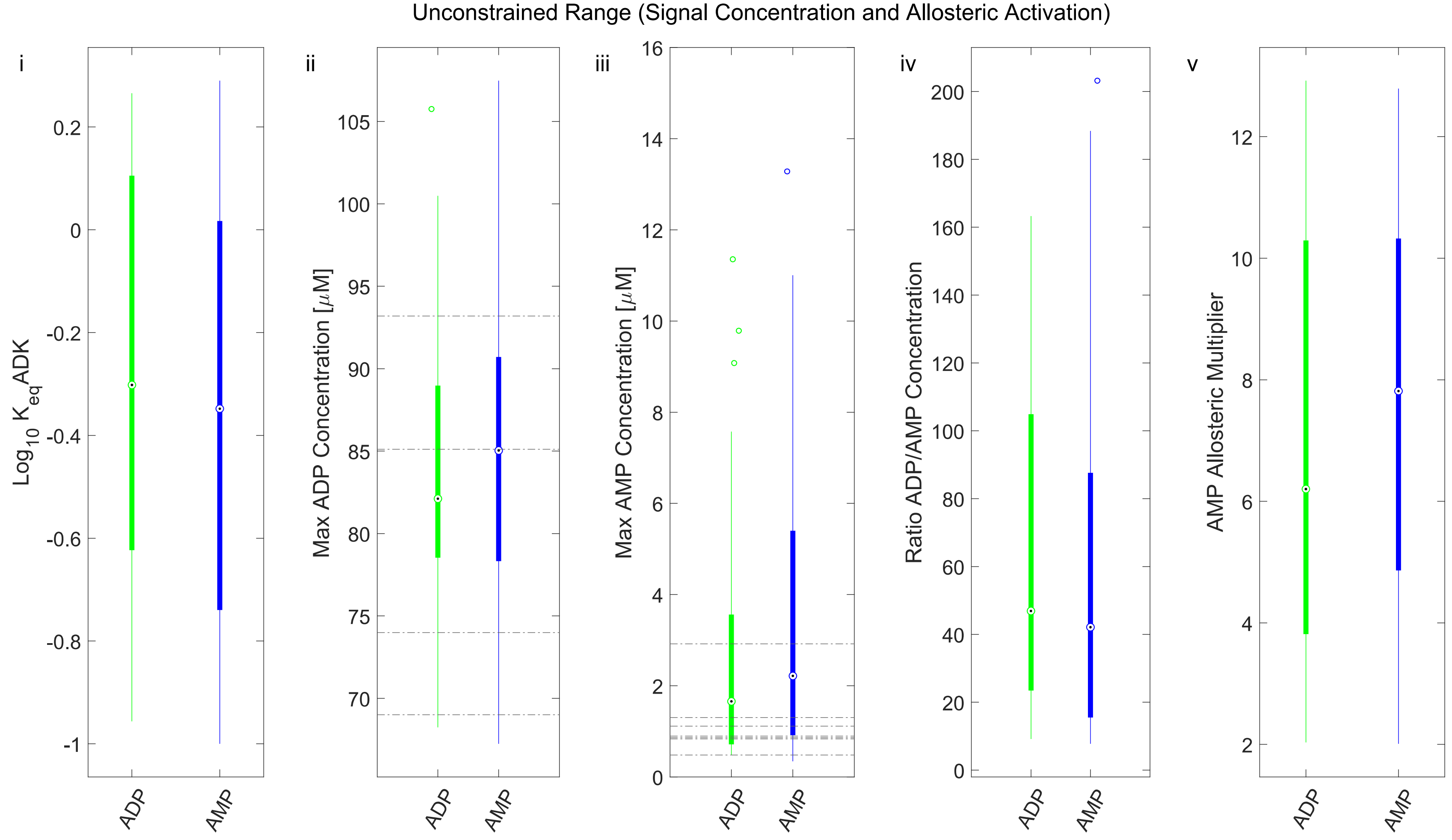


**Figure S1C**. Effects of $K_{eqADK}$, AMP and ADP concentrations, and allostery on AMP- versus ADP-dominant control of AMPK activity in the Unconstrained MPSA. Panels from left to right: i) Boxplots summarize the $K_{eq}$ADK values. ii) and iii) Boxplots summarize the maximum predicted concentrations of ADP (ii) and AMP (iii) during simulated exercise. The dashed-grey horizontal lines represent the reported maximum levels of ADP or AMP in moderate-intensity exercise studies. iv) Boxplots summarize the ratios of the maximum concentrations of ADP and AMP during simulated exercise. v) Boxplots summarize the magnitudes of allosteric activation of the AMP-p-AMPK complex.

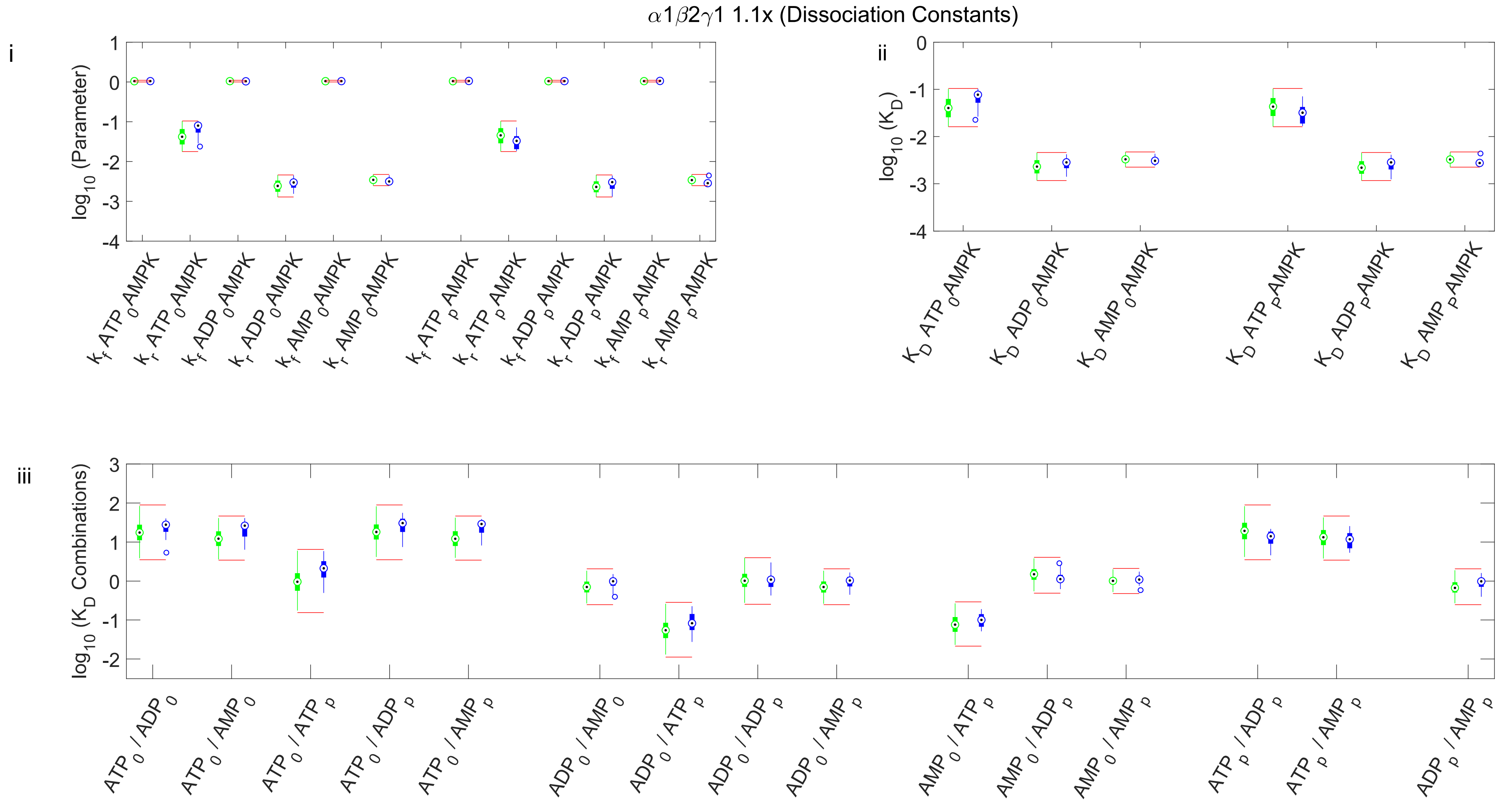


**Figure S2A**. Effects of AXP-(p)-AMPK binding constants on AMP- versus ADP-dominant control of AMPK activity in the **α1β2γ1 $K_D$ MPSA**. *i)* Boxplots summarize the individual parameter values representing the forward ($k_f$) and reverse ($k_r$) rate constants. *ii)* Boxplots summarize the dissociation constants ($K_D$) for each AXP-(p)-AMPK reaction, calculated as $k_r/k_f$ from each simulation. *iii)* Boxplots summarize the ratios of all possible pairs of $K_D$ values.

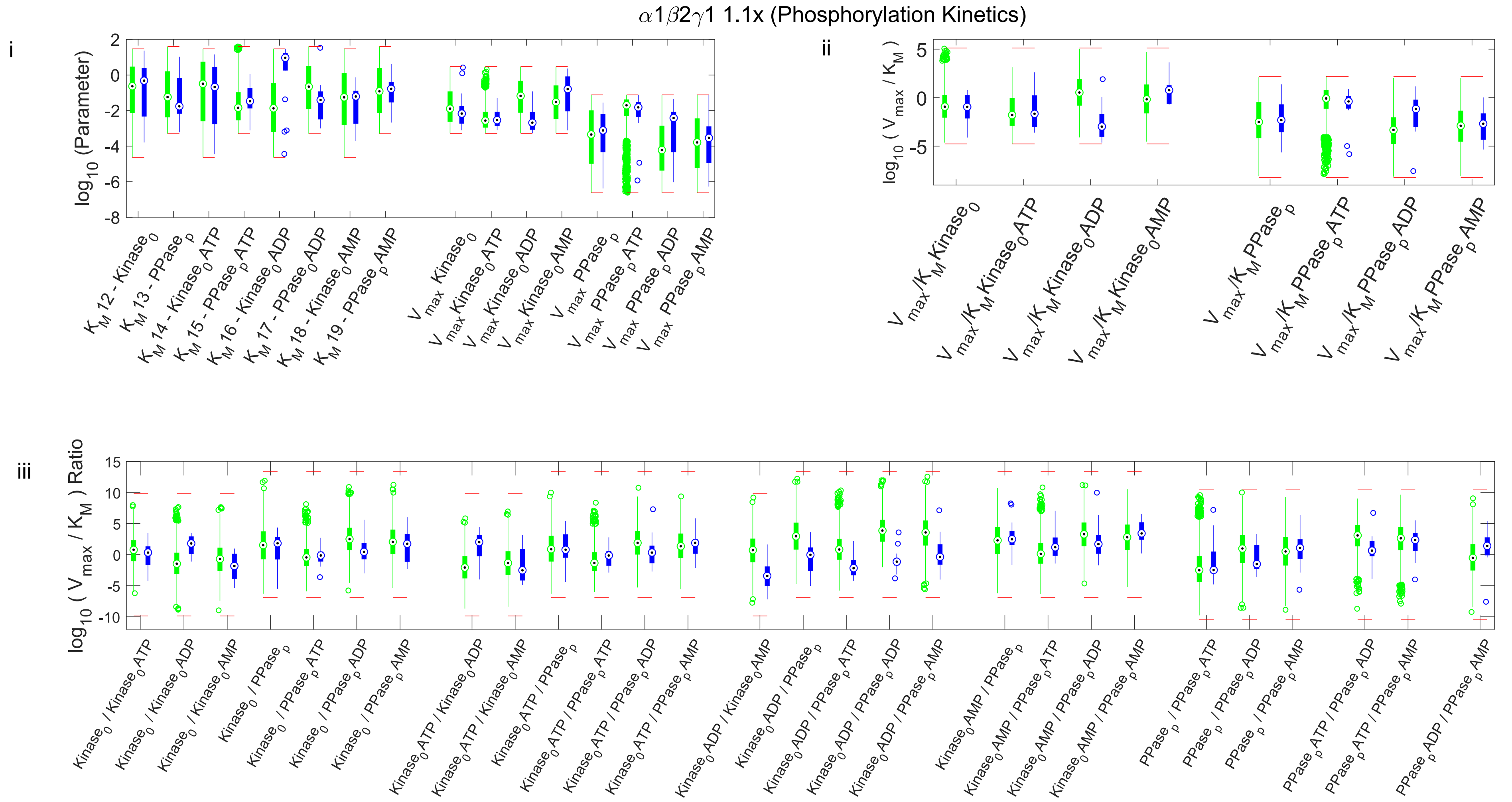


**Figure S2B**. Effects of parameters determining the enhancement of AMPK phosphorylation on AMP- versus ADP-dominant control of AMPK activity in the α1β2γ1 $K_D$ MPSA. *i)* Boxplots summarize the Michaelis constants ($K_M$) and maximum enzyme velocities ($V_{max}$) for the AMPK kinase and phosphatase. *ii)* Boxplots summarize the catalytic efficiencies of the AMPK kinase and phosphatase, which was calculated as $V_{max}/K_M$ for each simulation. *Bottom panel:* Boxplots summarize the compound ratios of kinase and phosphatase catalytic efficiencies.

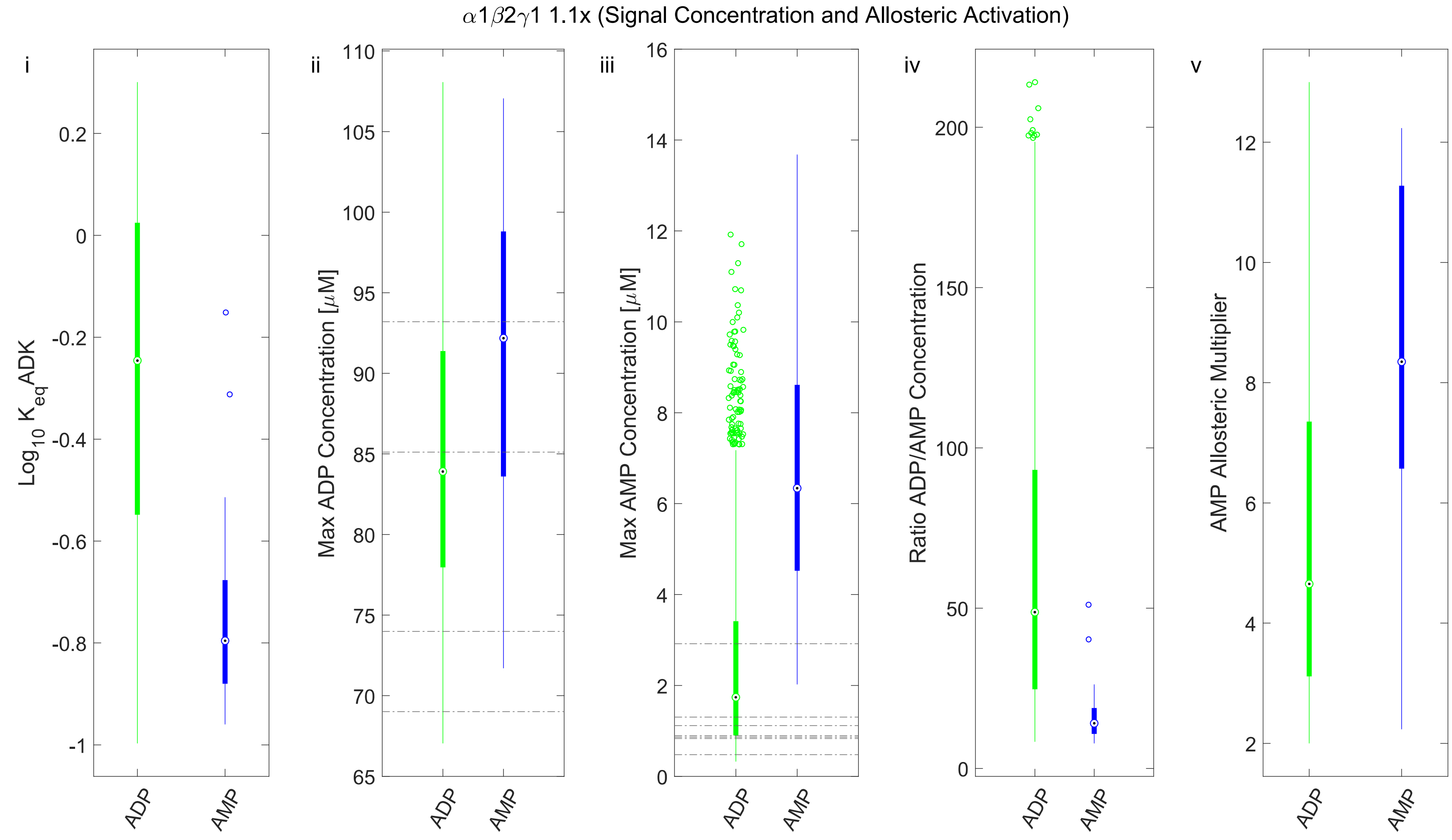


**Figure S2C**. Effects of $K_{eqADK}$, AMP and ADP concentrations, and allostery on AMP- versus ADP-dominant control of AMPK activity in the α1β2γ1 $K_D$ MPSA. Panels from left to right: *i)* Boxplots summarize the $K_{eq}$ADK values. *ii)* and *iii)* Boxplots summarize the maximum predicted concentrations of ADP (*ii*) and AMP (*iii*) during simulated exercise. The dashed-grey horizontal lines represent the reported maximum levels of ADP or AMP in moderate-intensity exercise studies. *iv)* Boxplots summarize the ratios of the maximum concentrations of ADP and AMP during simulated exercise. *v)* Boxplots summarize the magnitudes of allosteric activation of the AMP-p-AMPK complex.

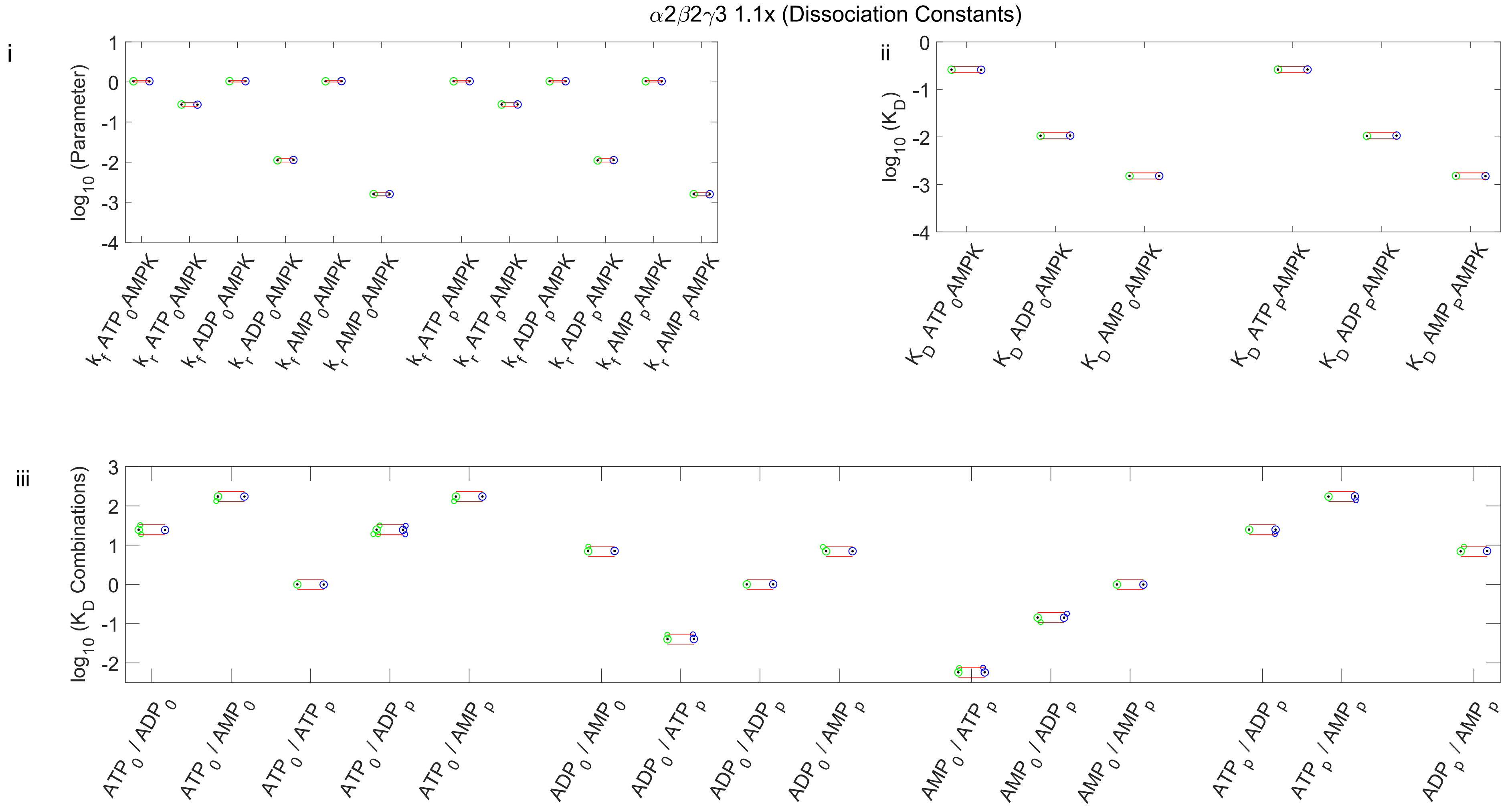


**Figure S3A**. Effects of AXP-(p)-AMPK binding constants on AMP- versus ADP-dominant control of AMPK activity in the **α2β2γ3 $K_D$ MPSA**. *i)* Boxplots summarize the individual parameter values representing the forward ($k_f$) and reverse ($k_r$) rate constants. *ii)* Boxplots summarize the dissociation constants ($K_D$) for each AXP-(p)-AMPK reaction, calculated as $k_r/k_f$ from each simulation. *iii)* Boxplots summarize the ratios of all possible pairs of $K_D$ values.

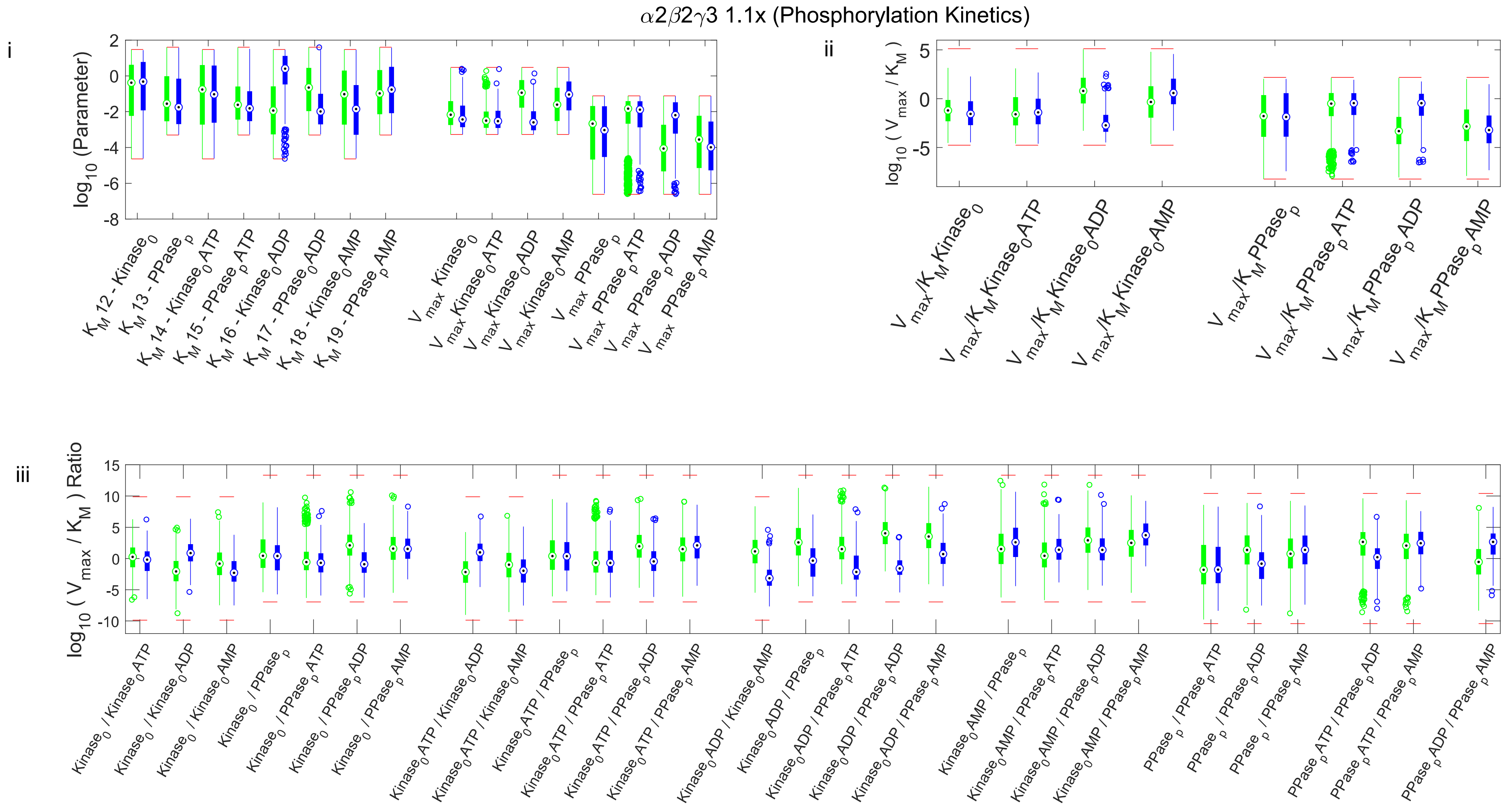


**Figure S3B**. Effects of parameters determining the enhancement of AMPK phosphorylation on AMP- versus ADP-dominant control of AMPK activity in the α2β2γ3 $K_D$ MPSA. i) Boxplots summarize the Michaelis constants ($K_M$) and maximum enzyme velocities ($V_{max}$) for the AMPK kinase and phosphatase. ii) Boxplots summarize the catalytic efficiencies of the AMPK kinase and phosphatase, which was calculated as $V_{max}/K_M$ for each simulation. iii) Boxplots summarize the compound ratios of kinase and phosphatase catalytic efficiencies.

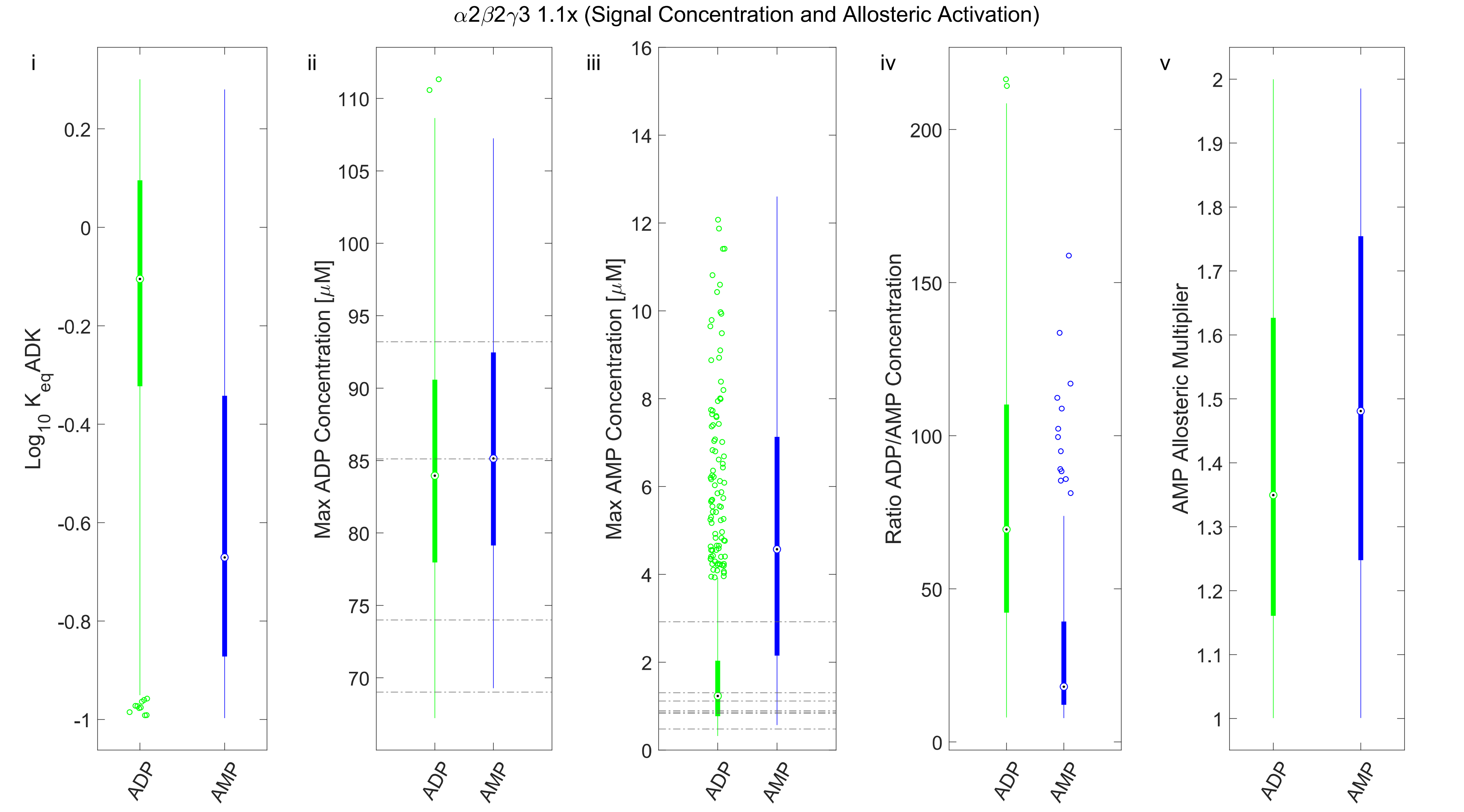


**Figure S3C**. Effects of $K_{eqADK}$, AMP and ADP concentrations, and allostery on AMP- versus ADP-dominant control of AMPK activity in the α2β2γ3 MPSA. Panels from left to right: i) Boxplots summarize the $K_{eq}$ADK values. ii) and iii) Boxplots summarize the maximum predicted concentrations of ADP (ii) and AMP (iii) during simulated exercise. The dashed-grey horizontal lines represent the reported maximum levels of ADP or AMP in moderate-intensity exercise studies. iv) Boxplots summarize the ratios of the maximum concentrations of ADP and AMP during simulated exercise. v) Boxplots summarize the magnitudes of allosteric activation of the AMP-p-AMPK complex.

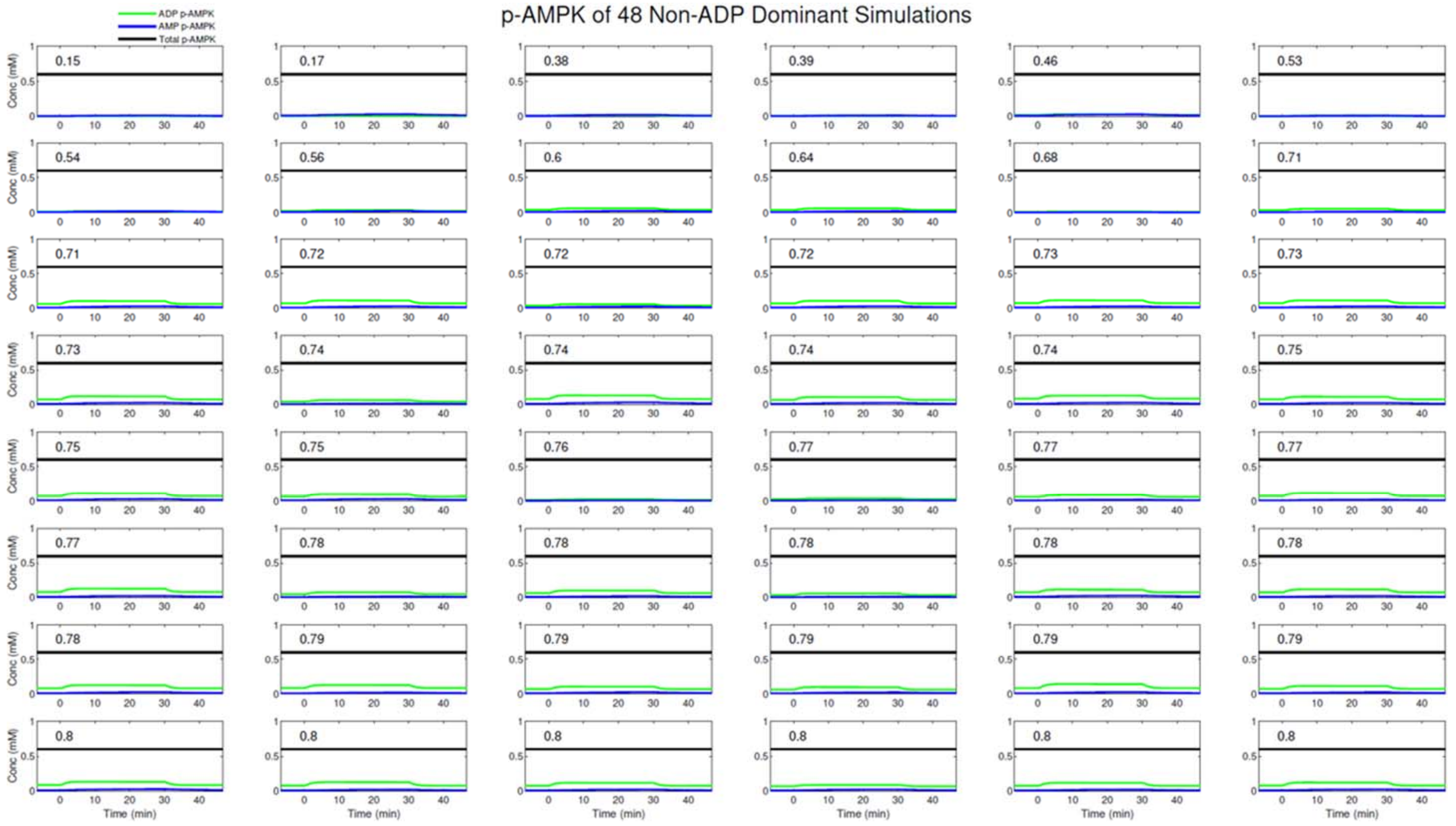


**Figure S4**. Model-predicted time courses of AMP-p-AMPK and ADP-p-AMPK in the 48 models remaining after the elimination process (see Fig. 4 of the main text). The blue, green, and black lines represent the levels of AMP-p-AMPK, ADP-p-AMPK and total AMPK, respectively. The inset numbers represent the corresponding fraction of ADP control.
